# Postmarketing commitments for novel drugs and biologics approved by the US Food and Drug Administration: a cross-sectional analysis

**DOI:** 10.1101/447185

**Authors:** Joshua D Wallach, Anita T Luxkaranayagam, Sanket S Dhruva, Jennifer E Miller, Joseph S Ross

**Affiliations:** Department of Environmental Health Sciences, Yale School of Public Health, 60 College Street, New Haven, CT, 06510, USA; Collaboration for Research Integrity and Transparency (CRIT), Yale University, 157 Church Street, 17^th^ Floor, Suite 1, New Haven, CT, 06510, USA; Center for Outcomes Research and Evaluation (CORE), Yale-New Haven Hospital, 1 Church Street, Suite 200, New Haven, CT, 06510, USA; University of Connecticut, Storrs, CT, 06269, USA; Department of Medicine, University of California, San Francisco School of Medicine, San Francisco, USA; Section of Cardiology, San Francisco Veterans Affairs Health Care System, San Francisco, CA, 4150 Clement St, Building 203, 2^nd^ Floor Cardiology, San Francisco, CA, 94121, USA; Section of General Medicine, Department of Internal Medicine, Yale School of Medicine, P.O. Box 208093, New Haven, CT, 06520-8093, USA; National Clinician Scholars Program, Department of Internal Medicine, Yale School of Medicine, New Haven CT, USA; Department of Health Policy and Management, Yale School of Public Health, 60 College Street, New Haven, CT, 06510, USA

**Author notes:** **Corresponding author**: Joshua D Wallach, MS, PhD Department of Environmental Health Sciences, Yale School of Public Health Collaboration for Research Integrity and Transparency (CRIT), Yale Law School 60 College Street, 4^th^ Floor, Room 411, New Haven, CT, 06510, USA.

**Keywords:** Postmarketing commitments, postmarketing requirements, FDA, lifecycle evaluation, pharmaceutical regulation

## Abstract

**Background:** Postmarketing commitments are clinical studies that drug sponsors agree to conduct at the time of FDA approval, but which are not required by statute or regulation. The objective of this study was to determine the characteristics, completion, and dissemination of postmarketing commitments agreed upon by sponsors at first FDA approval.

**Methods:** We performed a cross-sectional analysis of postmarketing commitments for new drugs and biologics approved 2009-2012. Using public FDA documents, ClinicalTrials.gov, and Scopus, we determined postmarketing commitments and their characteristics known at the time of FDA approval; number of postmarketing commitments subject to reporting requirements, for which FDA is required to make study status information available to the public (“506B studies”), and their statuses; and rates of registration and results reporting on ClinicalTrials.gov and publication in peer-reviewed journals for all clinical trials, with follow-up through July 2018.

**Results:** Among 110 novel drugs and biologics approved by the FDA between 2009-2012, 61 (55.5%) had at least one postmarketing commitment at the time of first approval. Of 331 total postmarketing commitments, 271 (81.9%) were non-human subjects research, predominantly chemistry, manufacturing, and controls studies; 49 (14.8%) were clinical trials (33 new and 16 ongoing trials for which follow-up results would be reported). Study descriptions for the new clinical trials often lacked information to establish study design features. Of the 89 (26.9%) 506B studies subject to public reporting requirements, of which 42 were clinical trials, 59 (66.3%) did not have an up-to-date status provided by FDA. Nearly all new clinical trials (28 of 31, 90.3%) were registered on ClinicalTrials.gov; of the 23 registered trials that were completed or terminated, 22 (95.7%) had reported results. Only half (14 of 29, 48.3%) of completed or terminated clinical trials, registered or unregistered, were published in peer-reviewed journals. Conclusions: The majority of postmarketing commitments agreed to by sponsors at the time of FDA approval for novel drugs and biologics approved between 2009-2012 were chemistry, manufacturing, and controls studies. While only 15% were clinical trials, these trials were nearly always registered with reported results on ClinicalTrials.gov. However, despite FDA public reporting requirements, up-to-date study status information was often unavailable for 506B studies.

## Background

Under the US Food and Drug Administration’s (FDA) lifecycle evaluation process, it is assumed that the benefit-risk balance of drugs and biologics will continue to be monitored after approval.[1, 2] Although FDA currently has four authorities that can be used to require New Drug Application (NDA) sponsors (generally pharmaceutical companies) to conduct studies in the postmarket setting (i.e., “postmarketing requirements”, **Box 1**),[3–5] additional clinical evidence, including long-term drug effectiveness data, can be generated through “postmarketing commitments”, which are “studies or clinical trials that a sponsor has agreed to conduct, but that are not required by a statute or regulations”.[5]

Prior to 2008, the term “postmarketing commitment” referred to all required, agreed-upon, and voluntary studies conducted by sponsors after FDA drug approval (**Box 1**).[5] Although FDA had the authority to require certain studies after approval, approximately 90% of all postmarketing studies between 1990 and 2004 were agreed-upon commitments.[6] However, once the FDA Amendments Act (FDAAA) went into effect on March 25, 2008, FDA began to distinguish between legally required studies and clinical trials (i.e., postmarketing requirements) and those that sponsors agreed to conduct but are not required (i.e., postmarketing commitments) (**Box 1**). While postmarketing commitments are not formally mandated by FDA, certain postmarketing commitments, including clinical studies, are subject to reporting requirements under section 506B (“506B studies”) of the Federal Food, Drug, and Cosmetic Act (**Box 2**). For 506B studies, sponsors must annually report to the FDA the status of postmarketing commitments, and the FDA must publicly report on the status of these commitments.[7]

Prior studies have focused exclusively on the characteristics,[8] completion,[9] and dissemination of postmarketing requirements,[8, 10, 11] but little is known about postmarketing commitments after the post-FDAAA changes. Given that postmarketing commitments may be a potentially important source of information about drug safety and effectiveness after market approval, we characterized the postmarketing commitments for all novel drugs and biologics approved between 2009 and 2012 using publicly available data sources, including their status and study characteristics, and for clinical trial ClinicalTrials.gov, as well as publication in peer reviewed journals.

### Box 1.

#### The history of US Food and Drug Administration’s postmarketing commitments and requirements

| **Box 1. The history of US Food and Drug Administration’s postmarketing commitments and requirements** |  |
| --- | --- |
| **Before FDAAA<sup>a</sup>** | **After FDAAA** |
| Required, agreed upon, and voluntary studies were classified as “*postmarketing commitments*” | Required studies were classified as “*postmarketing requirements*” and agreed upon and voluntary studies were classified as “*postmarketing commitments*” |
| **Statutory or regulatory authorities used to require postmarketing studies** |  |
| - *Animal Efficacy Rule*: Novel drugs can be approved when human efficacy studies and field trials may not be ethical and/or feasible, but FDA may require postmarket studies in humans<br>- *Pediatric Research Equity Act*: FDA can approve novel drugs for use in adults without corresponding studies for the same indication in the relevant pediatric population, but FDA can include deferred pediatric studies or clinical trials as postmarketing requirements<br>- *Accelerated Approval*: To expedite the approval of novel drugs that treat serious diseases and that fill unmet medical needs on the basis of surrogate or intermediate endpoints “reasonably likely” to predict clinical benefit,[12] FDA has the authority to require postmarket studies to confirm efficacy | - *Animal Efficacy Rule*<br>- *Pediatric Research Equity Act*<br>- *Accelerated Approval*<br>- *Food and Drug Administration Amendments Act (FDAAA) Section 505(o)(3)*: To provide additional information for novel therapeutics approved under section 505 of FDAAA or section 351 of the Public Health Services Act, FDA can require postmarket studies that assess known serious risks, signs of serious risks, or unexpected serious risks related to the use of a novel drug |
| FDAAA = Food and Drug Administration Amendments Act<br><sup>a</sup>Section 901 of FDAAA went into effect on March 25, 2008 |  |

### Box 2.

#### Postmarketing commitment reporting requirements

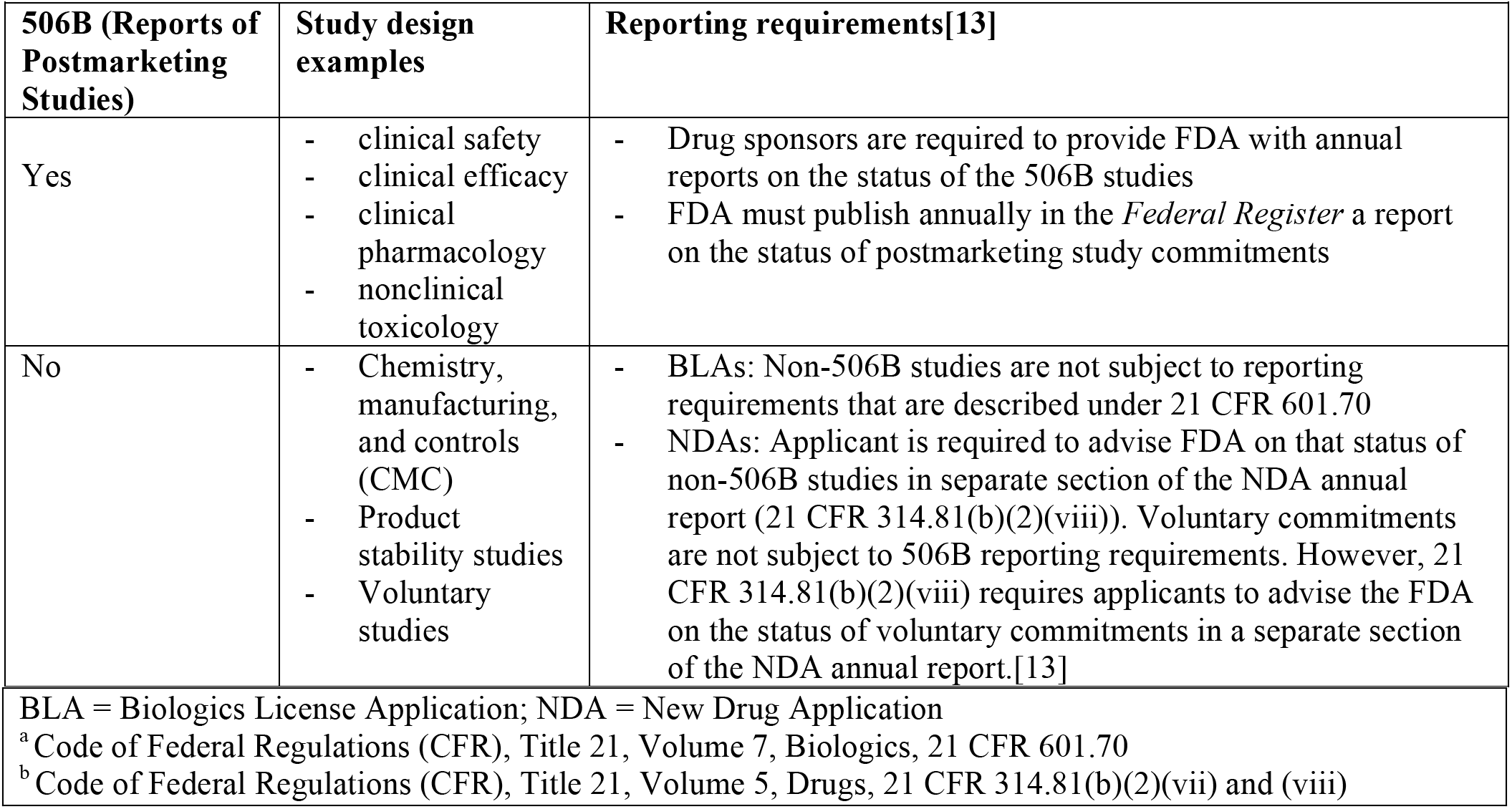

| **Box 2. Postmarketing commitment reporting requirements** |  |  |
| --- | --- | --- |
| **506B (Reports of Postmarketing Studies)** | **Study design examples** | **Reporting requirements[13]** |
| Yes | - clinical safety<br>- clinical efficacy<br>- clinical pharmacology<br>- nonclinical toxicology | - Drug sponsors are required to provide FDA with annual reports on the status of the 506B studies<br>- FDA must publish annually in the *Federal Register* a report on the status of postmarketing study commitments |
| No | - Chemistry, manufacturing, and controls (CMC)<br>- Product stability studies<br>- Voluntary studies | - BLAs: Non-506B studies are not subject to reporting requirements that are described under 21 CFR 601.70<br>- NDAs: Applicant is required to advise FDA on that status of non-506B studies in separate section of the NDA annual report (21 CFR 314.81(b)(2)(viii)). Voluntary commitments are not subject to 506B reporting requirements. However, 21 CFR 314.81(b)(2)(viii) requires applicants to advise the FDA on the status of voluntary commitments in a separate section of the NDA annual report.[13] |

BLA = Biologics License Application; NDA = New Drug Application
<sup>a</sup> Code of Federal Regulations (CFR), Title 21, Volume 7, Biologics, 21 CFR 601.70
<sup>b</sup> Code of Federal Regulations (CFR), Title 21, Volume 5, Drugs, 21 CFR 314.81(b)(2)(vii) and (viii)

## Methods

### Study design and sample

As in prior work focused on postmarketing requirements,[8] we used the publicly available Drugs@FDA database to identify and categorize all novel drug and biologic license applications (excluding generic drugs, reformulations, and combination therapies of non-novel therapeutic agents) first approved between January 1, 2009, and December 31, 2012.[14, 15] As previously described,[14] we characterized each new drug and biologic by date of approval, as a pharmacologic entity (small molecule) or biologic, and by orphan status; determined the first-approved indication for each new drug and biologic and whether applications were designated by FDA for priority review status.[16] We then used the World Health Organization’s anatomic therapeutic classification system to categorize each indication and grouped each indication into one of six therapeutic areas.[17]

### Identifying Postmarketing Commitments and Postmarketing Commitments Features

One author (ATL) identified all postmarketing commitments and dates that the FDA sets for important milestones (i.e., final protocol submission, trial completion, and final report submission), which are outlined in the approval letters hyperlinked in the Drugs@FDA database. These letters include a brief description of the study type and outline whether commitments are subject to certain reporting requirements (**Box 2**). In particular, section 506B of the Food, Drug, and Cosmetic Act (506B studies), 21 Code of Federal Regulations (CFR) 314.81(b)(2)(vii), and CFR 601.70,[4, 7] require drug sponsors to report annually on the status of certain postmarketing commitments. Additionally, for 506B studies, the FDA must publish annually in the *Federal Register* a report on the status of postmarketing commitments.[4] We then classified each postmarketing commitment into one of four study categories about the type of study required (**Box 3**), calculated the length of each postmarketing commitment study description (word count), and abstracted key study design characteristics using a previously described approach. [8]

#### Box 3.

##### Postmarking commitment categorization

###### New clinical trials

Postmarketing commitments that outline *new* clinical trials, including randomized and non-randomized clinical trials evaluating efficacy or “efficacy and safety”. This includes “clinical trials in which the primary endpoint is related to further defining efficacy, designed to: evaluate long-term effectiveness or duration of response, evaluate efficacy using a withdrawal design, evaluate efficacy in a subgroup.”[4]

###### Complete or submit results from ongoing clinical trials

Instead of requesting *new* clinical trials, these postmarketing commitments call for the completion and submission of the results from ongoing clinical trials.

###### Observational studies, analyze/follow-up from clinical studies, and other flexible commitments

Postmarketing commitments that outline longer follow-up or new analyses of data from existing trials or studies; submission of a final report for ongoing case-control, cross-sectional, or retrospective cohort studies.

###### Other studies

Manufacturing, stability, and immunogenicity studies that do not have a primary safety endpoint; pharmacoepidemiologic studies; pharmacokinetic and/or pharmacodynamics trials; and chemistry, manufacturing, and controls study commitments that sponsors have agreed with the FDA to conduct (CMC commitments).

### Status of Postmarket Studies

We used Postmarketing Study and Clinical Trial Requirements and Commitments Database Files to determine the status (i.e., Pending, Ongoing, Delayed, Terminated, Submitted, Fulfilled, or Released. **Additional file 1: supplementary Box 1**) of commitments specifically classified as subject to reporting requirements under 506B (**Box 2**). [18] We downloaded the most recent Postmarketing Study and Clinical Trial Requirements and Commitments Database file on August 2, 2018 (representing data updated by FDA as of July 24, 2018). Previous Postmarketing Study and Clinical Trial Requirements and Commitments Database Files were located using the FDA.gov Archive. When archived databases with the final statuses were unavailable, we recorded the most recent status and date for each postmarketing commitment (e.g., “last available status: *Delayed*, October 31, 2010”). We then performed additional Google searches using the terms “postmarketing requirement”, “postmarketing commitment”, “PMR”, or “PMC” in combination with the manufacturer or drug brand name to determine whether manufacturers were publicly sharing their own information about postmarketing commitments (e.g., “Pfizer PMC” or “Pfizer postmarketing commitment”). For each Google search, we screened the first 100 results. Lastly, we reviewed all supplemental letters on the Drugs@FDA database to determine whether they included information regarding the fulfillment of postmarketing commitments. Abstractions were performed by one reviewer (ATL) and consistency and accuracy were verified by a second reviewer (JDW).

### Trial Registration and Results Reporting on ClinicalTrials.gov and Peer-Reviewed Publication

For all new clinical trials and all commitments that call for the completion and submission of the results from ‘ongoing’ clinical trials (**Additional file 1: supplementary Boxes 2 and 3**), we determined study registration and results reporting on ClinicalTrials.gov, as previously described.[8] If identified, for each registered clinical trial, one reviewer (JDW) abstracted study characteristics from the ClinicalTrials.gov registration. The primary outcome providing the highest level of evidence was recorded. For instance, for trials with multiple primary efficacy outcomes, we considered clinical outcomes the highest level, followed by clinical scales and surrogate markers. A third reviewer (SSD) repeated all searches for trials that were determined to be unregistered, and uncertainties were discussed with the senior investigator (JSR).

For all clinical trials with a *Completed* or *Terminated* status on ClinicalTrials.gov for which results reporting would be expected, we abstracted whether any study results were reported and/or corresponding articles were published. For *Completed* or *Terminated* trials without reported results, we determined whether the date of final data collection for the prespecified primary outcome measure(s) (primary completion date) was within 12 months of the follow-up date (July 2018). According to the Final Rule for Clinical Trials Registration and Results Information Submission (“Final Rule”), submission of final results information is required “not later than 1 year after the completion date.”[19] For all clinical trials without publications listed on ClinicalTrials.gov and all unregistered clinical trials classified as *Submitted, Fulfilled, Released*, or unclear (i.e., no status available) according to FDA or drug sponsor data, one reviewer (JDW) used a systematic two-step search strategy to locate publications, as has been done in prior research.[8, 20] A third reviewer (SSD) repeated all searches for postmarketing commitments that were determined to be unpublished.

### Statistical Analysis

We used descriptive statistics to characterize the new drugs and biologics and postmarketing commitments. Analyses were performed using R (version 3.2.3; The R Project for Statistical Computing).

## Results

### Characteristics of New Drugs and Biologics

Between 2009 and 2012, FDA approved 110 new drugs and biologics for 120 indications. Of these, 49 (44.5%) did not have any postmarketing commitments at the time of first approval. Among the 61 drugs and biologics for 68 total indications in the final sample (**Table 1**), 39 (63.9%) were drugs, 22 (36.1%) biologics; 19 (31.2%) were indicated for the treatment of cancer or hematologic disease; and 21 (34.4%) received priority review. There were 7 (11.5%) drugs and biologics that received accelerated approval and 14 (23.0%) that were designated as orphan products.

**Table 1.**
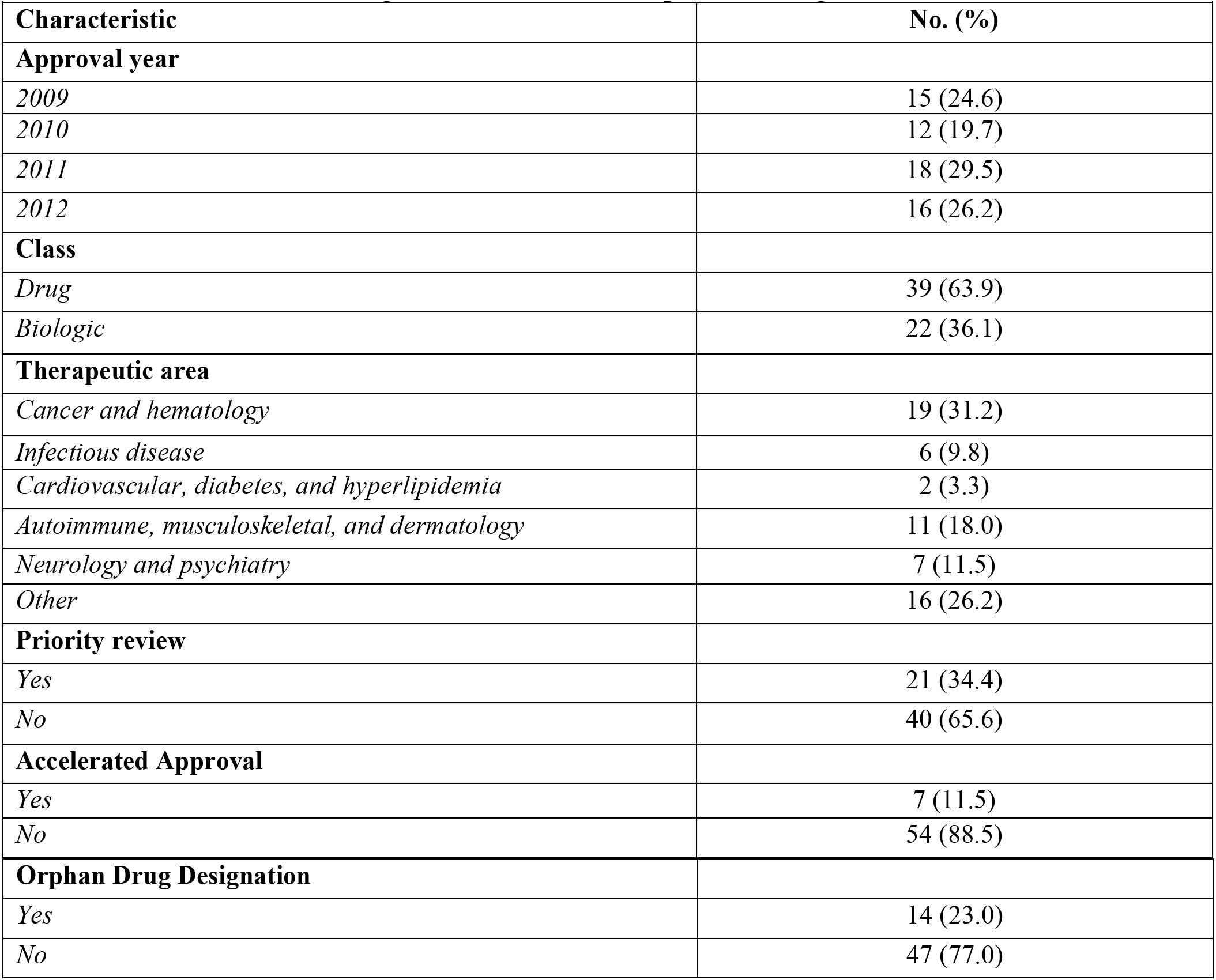
Characteristics of 61 new drugs and biologics approved by the US Food and Drug Administration from 2009 through 2012 with at least one postmarketing commitment

|  |  |
| --- | --- |
| **Characteristic** | **No. (%)** |
| **Approval year** |  |
| *2009* | 15 (24.6) |
| *2010* | 12 (19.7) |
| *2011* | 18 (29.5) |
| *2012* | 16 (26.2) |
| **Class** |  |
| *Drug* | 39 (63.9) |
| *Biologic* | 22 (36.1) |
| **Therapeutic area** |  |
| *Cancer and hematology* | 19 (31.2) |
| *Infectious disease* | 6 (9.8) |
| *Cardiovascular, diabetes, and hyperlipidemia* | 2 (3.3) |
| *Autoimmune, musculoskeletal, and dermatology* | 11 (18.0) |
| *Neurology and psychiatry* | 7 (11.5) |
| *Other* | 16 (26.2) |
| **Priority review** |  |
| *Yes* | 21 (34.4) |
| *No* | 40 (65.6) |
| **Accelerated Approval** |  |
| *Yes* | 7 (11.5) |
| *No* | 54 (88.5) |

| Orphan Drug Designation |  |
| --- | --- |
| *Yes* | 14 (23.0) |
| *No* | 47 (77.0) |

### Postmarketing Commitments, 2009-2012

The FDA approval letters for these 61 drugs and biologics described 331 separate postmarketing commitments. The median number of commitments per approval letter was 3 (interquartile range [IQR], 1 to 7). The majority of the postmarketing commitments (271 of 331 (81.9%)) were ‘Other studies’, including chemistry, manufacturing, and controls (CMC) studies (**Additional file 1: supplementary box 3**). Of the 49 (14.8%) commitments outlining a clinical trial, 33 described new clinical trials and 16 called for the submission of final reports or data from ongoing trials. Just over one-quarter (89 (26.9%)) of the postmarketing commitments were subject to 506B reporting requirements (**Table 2**).

**Table 2.**
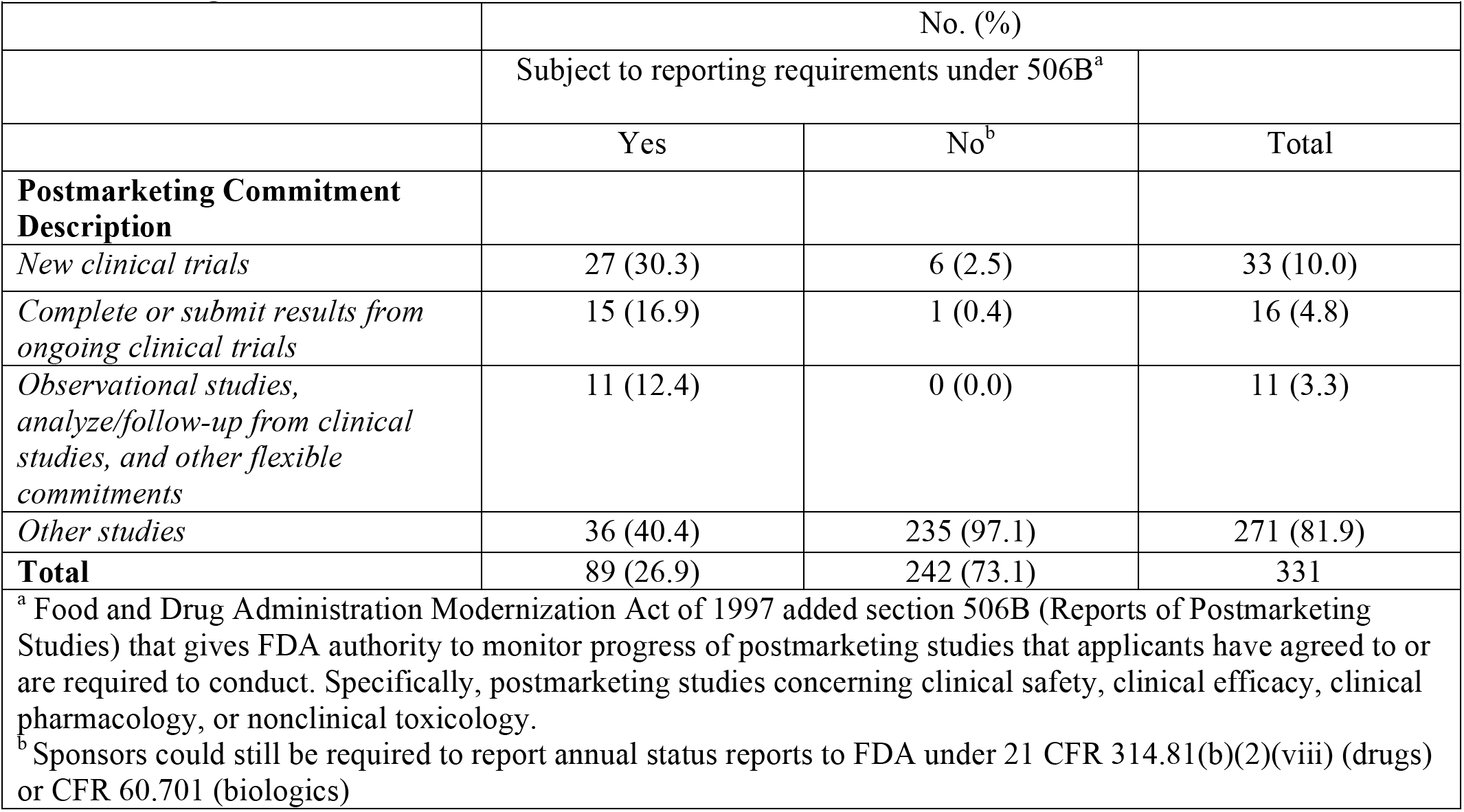
Categories of postmarketing commitments for novel drugs and biologics approved by the US Food and Drug Administration between 2009 and 2012

|  |  |  |  |
| --- | --- | --- | --- |
|  | No. (%) |  |  |
|  | Subject to reporting requirements under 506B<sup>a</sup> |  |  |
|  | Yes | No<sup>b</sup> | Total |
| **Postmarketing Commitment Description** |  |  |  |
| *New clinical trials* | 27 (30.3) | 6 (2.5) | 33 (10.0) |
| *Complete or submit results from ongoing clinical trials* | 15 (16.9) | 1 (0.4) | 16 (4.8) |
| *Observational studies, analyze/follow-up from clinical studies, and other flexible commitments* | 11 (12.4) | 0 (0.0) | 11 (3.3) |
| *Other studies* | 36 (40.4) | 235 (97.1) | 271 (81.9) |
| **Total** | 89 (26.9) | 242 (73.1) | 331 |
| <sup>a</sup> Food and Drug Administration Modernization Act of 1997 added section 506B (Reports of Postmarketing Studies) that gives FDA authority to monitor progress of postmarketing studies that applicants have agreed to or are required to conduct. Specifically, postmarketing studies concerning clinical safety, clinical efficacy, clinical pharmacology, or nonclinical toxicology.<br><sup>b</sup> Sponsors could still be required to report annual status reports to FDA under 21 CFR 314.81(b)(2)(viii) (drugs) or CFR 60.701 (biologics) |  |  |  |

Among the 33 postmarketing commitments for new clinical trials, the median number of words used to describe the study in publicly available documents was 42 (IQR, 29 to 63), thus providing limited information. Using only FDA approval letters, there was not enough information to establish use of randomization, allocation, comparator type, outcome, and number of patients to be enrolled for 13 (39.4%), 20 (60.6%), 27 (81.8%), and 30 (90.9%) of the 33 new clinical trials, respectively (**Additional file 1: supplementary table 1**). In contrast, all 16 postmarketing commitments calling for the submission of final reports or data for ongoing trials included either a trial name or identifier.

Among the 89 postmarketing commitments subject to 506B reporting requirements, 20 (22.5%) were classified as fulfilled according to FDA Postmarketing Study and Clinical Trial Requirements Database files. Over two-thirds (59 of 89 (66.3%)) of the 506B studies did not have enough information in the databases to determine an up to data status; 35 of these had no past or current status (**Additional file 1: upplementary table 2**).

Among the 27 postmarketing commitments for new clinical trials subject to 506B reporting requirements, 12 (44.4%) did not have an up-to-date status and 8 (29.7%) were classified as fulfilled. When all FDA supplemental letters and drug sponsor data were considered in addition to the databases, 40 (40 of 89, 44.9%) were classified as fulfilled and 38 (42.7%) did not have enough information to determine an up to date status (**Additional file 1: supplementary table 3**). Publicly available drug sponsor data were available for 28 506B postmarketing commitments.

### Registration and Study Characteristics of Clinical Trials

Among the 33 postmarketing commitments for new clinical trials, two did not have enough information in their postmarketing commitment descriptions to perform ClinicalTrials.gov searches. Of the 31 remaining commitments, 28 (90.3%) were registered on ClinicalTrials.gov (**Table 3**). The majority of the 28 registered clinical trials were randomized (26, 92.9%) with double or triple blinding (22, 78.6%) (**Table 4**). Eighteen (63.0%) trials were placebo controlled and 4 (14.3%) had an active comparator. The majority of trials included efficacy primary endpoints that were categorized as clinical scales (n=17; 60.7%) and clinical outcomes (n=5; 17.9%); 5 (17.9%) focused on surrogate markers of disease. The median study duration and estimated or actual sample size according to the ClinicalTrials.gov registrations were 1.6 months (IQR, 0.9 to 12.0) and 400 patients (IQR, 254 to 529) respectively.

**Table 3.** Registration, results reporting, and publication of postmarketing commitments of new drugs and iologics approved by the Food and Drug Administration between 2009 and 2012

| **Table 3. Registration, results reporting, and publication of postmarketing commitments of new drugs and biologics approved by the Food and Drug Administration between 2009 and 2012** |  |  |  |  |  |  |  |
| --- | --- | --- | --- | --- | --- | --- | --- |
|  | No. (%) |  |  |  |  |  |  |
|  | Registration |  | Results reporting |  | Publication<sup>a</sup> or results reporting |  |  |
| Category | Eligible for registration at ClinicalTrials.gov | Registered | Eligible for results reporting<sup>b</sup> | Results reported | Eligible for publication<sup>c</sup> | Published | Results reported or published |
| *New clinical trials* | 31 | 28 (90.3) | 23 | 22 (95.7) | 29 | 14 (48.3) | 22 (75.9) |
| *Complete or submit results from clinical trials* | 16 | 16 (100.0) | 15 | 15 (100.0) | 15 | 14 (93.3) | 15 (100.0) |
| Total | 49 | 44 (89.8) | 38 | 36 (94.7) | 44 | 28 (63.6) | 37 (84.1) |
| FDAAA = Food and Drug Administration Amendments Act<br><sup>a</sup>“Publication” indicates publication in the peer-reviewed literature.<br><sup>b</sup>Clinical studies classified as *Completed* or *Terminated* by ClinicalTrials.gov. Among four additional trials that classified as *Active, not recruiting* on ClinicalTrials.gov, one reported results and none were published. One article classified as *Completed* had an “Actual Primary Completion Data” of April 30, 2018. The final rule requires the submission of results information “not later than 1 year after the completion data (referred to as the ‘primary completion date’).”<br><sup>c</sup>The denominator included *Completed* and *Terminated* clinical trials registered on ClinicalTrials.gov. For registered and unregistered clinical trials, we used information provided by FDA or drug sponsors on the status of the postmarketing commitments. We searched for publications for clinical trials classified by the FDA as *Submitted*, *Fulfilled*, or *Released*. We also searched for publications for postmarketing commitments where the last status provided by the FDA was unclear (e.g., last available record: 2013, ongoing). |  |  |  |  |  |  |  |

**Table 4.** Study characteristics of clinical trials based on ClinicalTrials.gov data

|  |  |  |  |  |  |  |  |  |  |  |  |  |
| --- | --- | --- | --- | --- | --- | --- | --- | --- | --- | --- | --- | --- |
|  | No. (%) |  |  |  |  |  |  |  |  |  | Median (IQR)<sup>a</sup> |  |
|  | Allocation |  |  | Comparator |  |  | Endpoint |  |  |  | Estimated Sample Size | Duration (months) |
| Registered Postmarketing Requirements | Randomized | Double Blind | Open label | Placebo | Active | None | Surrogate Outcome | Clinical Outcome | Clinical Scale | Safety |  |  |
| New clinical trial (n=28) | 26 (92.9) | 22 (78.6) | 6 (21.4) | 18 (64.3) | 4 (14.3) | 6 (21.4) | 6 (21.4) | 5 (17.9) | 17 (60.7) | 0 (0.0) | 400 (254 to 529) | 1.6 (0.9 to 12.0) |
| Complete or submit results from clinical trials (16) | 10 (62.5) | 6 (37.5) | 10 (62.5) | 6 (37.5) | 4 (25.0) | 6 (37.5) | 10 (62.5) | 3 (18.8) | 0 (0.0) | 3 (18.8) | 237 (373 to 744) | 5.6 (5.2 to 12.6) |
| <sup>a</sup> Registered on ClinicalTrials.gov |  |  |  |  |  |  |  |  |  |  |  |  |

All 16 postmarketing commitments outlining the completion or submission of results from clinical trials were registered on ClinicalTrials.gov (**Table 3**). Of these, 10 (62.5%) were randomized, 6 (37.5%) were double or triple blind, 6 (37.5%) were placebo controlled, and 4 (25.0%) had an active comparator (**Table 4**). The majority of the trials (10, 62.5%) focused on surrogate outcomes. According to the ClinicalTrials.gov registrations, median study duration and estimated sample size were 5.6 months (IQR, 5.2 to 12.6) and 237 patients (IQR, 373 to 744), respectively.

### Results Reporting and Publication of Clinical Trials

Of the 23 postmarketing commitments for new trials classified as completed or terminated according to ClinicalTrials.gov, 22 (95.7%) had reported results (**Table 3**). Among the 29 registered or unregistered studies for which publication would be expected based on the most recent status provided by the FDA, drug sponsors, or on ClinicalTrials.gov, just under half were published in a peer reviewed journal (14 of 29 (48.3%)) and approximately three-quarters (22 of 29 (75.9%)) had either reported results or were published. The median time from FDA approval to reported results or publication of new trials was 65 months (IQR, 47 to 81).

Among the 22 trials with reported results or a publication with a “report submission” date provided in the FDA approval letters, 18 (81.8%) reported results on ClinicalTrials.gov after the FDA scheduled submission deadline (median 13 (IQR, 4 to 24) months afterwards). There were 4 (18.9%) that reported results ahead of schedule (median 13 months (IQR, 10 to 16)). All 15 (100%) postmarketing commitments outlining the completion or submission of results from an ongoing study reported results, and all but one were published (14, 93.3%).

## Discussion

Just over half of the new drugs and biologics approved by the FDA between 2009-2012 had at least one postmarketing commitment outlined at the time of approval. These studies, which sponsors agree to conduct, are not required by a statute or regulations, but may be a potentially important source of information about drug safety and effectiveness after market approval. However, the vast majority were not human subjects research intended to gather additional information about the safety, efficacy, or optimal use of drugs and biologics in patients and were instead focused on product quality control, an important component of safety. Instead, only 15% were clinical studies, less than one in ten new clinical trials. While these trials were nearly always registered with reported results on ClinicalTrials.gov, approximately one-half had not yet been published in peer-reviewed journals and the vast majority reported results on ClinicalTrials.gov after FDA scheduled submission deadlines. Postmarketing commitments may offer an opportunity for FDA to work with sponsors to generate important information about recently approved drugs and biologics after market approval.

Our study found that over 80% of the postmarketing commitments were for non-clinical studies, including chemistry, manufacturing, and controls study commitments. Prior to FDAAA, when the term “postmarketing commitment” referred to all required, agreed-upon, and voluntary studies conducted by sponsors after FDA drug approval, 74% of commitments were classified as clinical studies.[21] However, since 2008, FDA has distinguished between legally required studies and clinical trials (postmarketing requirements) and studies that “would not meet the statutory purposes” for postmarketing requirements (postmarketing commitments).[4] Therefore, the low proportion of clinical trials identified as postmarketing commitments in our sample may not be surprising, considering that confirmatory, safety, and pediatric clinical trials can be formally required by FDA under the accelerated approval, FDAAA, and PREA postmarketing requirement authorities.[8, 11] However, postmarketing commitments can still include clinical trials “in which the primary endpoint is related to further defining efficacy”,[4] and certain clinical efficacy, clinical pharmacology, or nonclinical toxicology studies are subject to reporting requirements by applicants and the FDA.

Using only FDA’s Postmarketing Study and Clinical Trial Requirements Database files, we were unable to identify an up-to-date status for 68% of the postmarketing commitments subject to FDA reporting requirements under 506B. Drug sponsors are required to provide the FDA with annual status reports for 506B studies of certain agreed-upon commitments, and FDA must publish annually in the *Federal Register* a report on the status of these postmarketing study commitments. We found that among the new clinical trials subject to 506B reporting requirements, just under half had an up-to-date status and less than one-third were classified as fulfilled. These findings are consistent with a previous study suggesting that postmarketing requirements often lack a publicly available up-to-date status.[8] However, we also found that the rates of registration and results reporting on ClinicalTrials.gov and publication in peer-reviewed journal among postmarketing commitment clinical trials were promising, which was similar to what has been previously observed among postmarketing requirements.[8]

Over 80% of the postmarketing commitments for new clinical trials reported results on ClinicalTrials.gov after FDA scheduled submission deadlines, which is higher than what has been previously observed among postmarketing requirements (68.1%).[8] While FDA and drug sponsors can revise the milestones outlined in the initial approval letters, these findings suggest that there are delays in study conduct, completion and reporting, which may lead to gaps in the understanding of drug and biologic safety and effectiveness.

Prior studies have focused exclusively on the purposes and transparency of postmarketing requirements.[8, 22–25] Although postmarketing commitments are primarily for non-clinical studies, our work suggests that some commitments also generate clinical evidence. In order to further support FDA’s lifecycle evaluation process, FDA and drug sponsors should continue to work closely to identify new clinical studies or other studies already being conducted, beyond the confirmatory and safety postmarketing requirement that can be required by FDA. However, greater transparency will be necessary to ensure that data from postmarketing commitments are able to inform care. For instance, longer and more detailed study descriptions will allow for the identification of specific study design characteristics, including endpoints, which are necessary to inform clinical practice, as well as more detailed explanations about the potential long-term knowledge gaps that are addressed. Furthermore, FDA could consider expanding its recent plans to add ClinicalTrials.gov identifiers to materials for future drug approvals [26, 27] to include postmarketing commitments, especially for those describing ongoing studies with trial identifiers. Similarly, ClinicalTrials.gov could include a variable specifying whether certain trials are postmarketing requirements or commitments. This will allow patients, clinicians, and researchers to locate and identify potential postmarketing studies and their results. In order to ensure that postmarketing commitment status are publicly identifiable, the FDA should keep all “fulfilled” and “released” 506B commitments on the Postmarketing Study and Clinical Trial Requirements Database, instead of removing them after 1 year of fulfillment or completion. Lastly, although postmarketing commitment trails were often registered with reported results, drug sponsors can play a key role in promoting the dissemination of postmarket evidence by ensuring that the results of all agreed-upon clinical trials are published in peer-reviewed journals.

### Limitations of this study

This study has some limitations. First, by limiting our study to new approvals between 2009 and 2012, we did not identify and classify all postmarketing commitments issued after FDAAA. However, by focusing on this time period, we allowed for at least four years for completion, results reporting, and publication of postmarketing commitments. In our sample, the median study duration among clinical trials was 1.6 months, and only one-quarter of clinical trials had durations longer than 12 months. Therefore, we are reassured that our study allowed for an adequate amount of follow-up tome for studies to be completed, reported, and published. Second, as previously discussed, our study was designed to rely on publicly available data sources, which made determining study design characteristics, ClinicalTrials.gov registrations, and corresponding publications difficult in some cases.[8] Although we attempted to be comprehensive by using numerous public data sources, we were unable to locate an up-to-date status for nearly half of the commitments. Therefore, it is possible that a number of commitments classified as “unclear” are actually “fulfilled” and “released” commitments, which are only displayed on the FDA’s online database for one year after the date of fulfillment or release. Third, we used the milestone dates outlined in the initial FDA approval letters. However, sponsors can submit revised schedules and FDA uses the original study schedule to determine study progress.[13] Lastly, we also did not account for the time that it might take to prepare and publish research. Although additional studies could be published at a later date, but were not published at the time of our search, we based our decisions on previous work by our group and others.[8, 28–32]

### Conclusions

Among 331 postmarketing commitments outlined in approval letters for new drugs and biologics, the vast majority were for chemistry, manufacturing, controls, and other non-clinical studies. Only 15% of postmarketing commitments were new or ongoing clinical trials. While nearly half of postmarketing commitments subject to mandatory reporting requirements under Section 506B did not have a clear up-to-date progress reported publicly, the majority of clinical trials were registered on ClinicalTrials.gov with reported results. However, only half of the clinical trials had corresponding publications in peer-reviewed journals. Opportunities may exist for FDA and drug sponsors to work together to identify additional postmarketing commitments that support FDA’s lifecycle evaluation process by generating information about the safety, efficacy, or optimal use of drugs and biologics in patients.

## List of abbreviations

CI: confidence interval
IQR: Interquartile range
FDA: Food and Drug Administration

## Declarations

### Ethics approval and consent to participate

This study used publicly available information and did not require ethics approval from the Y ale University School of Medicine Human Research Protection Program.

### Consent for publication

Not applicable

### Availability of data and material

The datasets used and/or analysed during the current study are available from the corresponding author on reasonable request.

### Competing interests

In the past 36 months, JDW received research support through the Meta Research Innovation Center at Stanford (METRICS) from the Laura and John Arnold Foundation. JSR received research support through Yale from Johnson and Johnson to develop methods of clinical trial data sharing, from Medtronic, Inc. and the Food and Drug Administration (FDA) to develop methods for postmarket surveillance of medical devices (U01FD004585), from the Centers of Medicare and Medicaid Services (CMS) to develop and maintain performance measures that are used for public reporting, from the FDA to establish a Center for Excellence in Regulatory Science and Innovation (CERSI) at Yale University and the Mayo Clinic (U01FD005938), from the Blue Cross Blue Shield Association to better understand medical technology evaluation, and from the Agency for Healthcare Research and Quality (R01HS022882). SSD receives support as a Scholar in the Yale University / Mayo Clinic FDA CERSI. JEM and JSR receive funding from the Laura and John Arnold Foundation in connection with the Good Pharma Scorecard.

### Funding

This project was conducted as part of the Collaboration for Research Integrity and Transparency (CRIT) at Yale, funded by the Laura and John Arnold Foundation, which supports JDW and JSR. During the time of this study, SSD was supported by the Robert Wood Johnson Foundation’s Clinical Scholars Program and the Department of Veterans Affairs. These funders played no role in the design of the study, analysis or interpretation of findings, or drafting the manuscript and did not review or approval the manuscript prior to submission. The authors assume full responsibility for the accuracy and completeness of the ideas presented.

### Authors’ contributions

JDW, SSD, and JSR were responsible for the conception and design of this work. JDW and ATL were responsible for the data abstraction. Statistical analyses were performed by JDW. JDW drafted the manuscript, which was revised by JSR. All authors participated in the analysis and interpretation of the data and critically revised the manuscript for important intellectual content. All authors approved the final version of the manuscript. JSR provided supervision.

### Consent for publication

Not applicable.

### Additional files

Additional file 1: Supplementary tables and boxes (.PDF 11 KB)

